# Histopathological characteristics of different parts of surgical specimens of UPJ stenosis

**DOI:** 10.1101/2022.09.13.507884

**Authors:** NaFeisha TuErdi, Junbo Bai, Kaifang Liu, Shuai Liu, Hongxing Xiong, Yu Gao, Jia Li

## Abstract

**Objective:** To observe the pathological changes of different anatomical sites in the specimens obtained from ureteropelvic junction obstruction (UPJO) caused by UPJ stenosis.

**Materials and Methods:** A total of 34 cases of UPJO were performed. The lesion of the ureteropelvic junction was visualized as the center, the 1.5cm renal pelvis segment was taken along the upper edge of the lesion segment (as the control group). Along the lower edge of the ureteral stricture, 1cm of ureteral stricture tissue was taken downward. Different dyeing methods were used to observe the tissue arrangement of different parts, the ratio of muscle to fiber tissue and the distribution of interstitial cells of Cajal(ICC) .

**Results:** The number of ICCs in the lesion segment was reduced or even absent, disordered arrangement of muscle tissue was seen, the fibrous tissue proliferated to varying degrees. The pathological changes were statistically different from those of normal segment and ureteral stricture segment.

**Conclusion:** The decrease of ICCs cells and the degree of tissue fibrosis are closely related to the pathogenesis and disease progression of UPJO caused by UPJ stenosis.

## INTRODUCTION

Congenital ureteropelvic junction obstruction (UPJO) results in an unobstructed flow of urine from the renal pelvis into the proximal ureter, which gradually causes dilatation of the collecting system. It is a kind of congenital urinary system malformation that may cause varying degrees of renal function damage if the obstruction is not intervened in time, and it is also the most common cause of neonatal hydronephrosis. It is more common in males and has a higher incidence of left side lesions, and the disease due to UPJ stenosis is more common. For a long time, the etiology of UPJ stenosis in UPJO has been the focus of researchers. Over the years, a large number of clinical studies have found that its pathogenesis is affected by a variety of factors. Usually, it can be divided into intraluminal factors (such as: ureteral scar hyperplasia, ureteral dysplasia) and extraluminal factors (compression of lower renal vessels or cords, congenital renal malformation, iatrogenic ureteral scar, fibroepithelial polyps, etc.)[1]. Some information about renal embryogenesis has been obtained, but its etiology remains obscure. There is no reliable and definitive theory to explain this frequent anomaly. At present, most scholars believe that these processes may be related to the structural characteristics of UPJ.

The obstruction is caused by an undynamic stenosis segment, this lesion may be related to the dysplasia of the renal pelvis, the failure of recanalization of the ureteral bud junction, the abnormal concentration of molecules in some signaling pathways during embryonic development, the abnormal innervation of the renal pelvis, and the impaired differentiation of smooth muscle. A possible association with some single gene mutations has also been proposed.[2]

A large number of studies have been conducted on the etiology of UPJO at home and abroad, but the reason why functional UPJO appears and will disappear without intervention in most cases is still not clear. The aim of this study is to compare the pathological changes and differences among different parts of the tissue by observing the pathological changes in the surgical specimens of children with UPJ stenosis who underwent surgical treatment to search for possible causes of treatment.

## MATERIAlS AND METHODS

### 2.1 Selection of patients and obtain the samples

A total of 92 children with hydronephrosis were admitted to our hospital from January 1, 2021 to November 1, 2021, aged from 15 days to 14 years, excluding children older than 7 years and younger than 3 years, with an average age of 5.3 years. Our inclusion criteria were: It meets the diagnostic criteria of UPJO in Pediatric Urology, and has the support of auxiliary examination results such as B-ultrasound, nuclear magnetic resonance hydrography, intravenous pyelography, urological contrast-enhanced ultrasound, and so on. Excludes cases caused by high openings, ectopic vascular compression, valves, polyps, and other factors, only cases caused by stenosis were considered. It conforms to the surgical indications of pyeloplasty in the expert consensus. All operations were performed by the same surgeon and assisted by the same team. The specimens of the ureteropelvic junction (UPJ) were obtained by disconnection pyeloplasty, and the informed consent for specimen processing was signed by the family members. The exclusion criteria were: Severe disorders of the heart, brain, lung, and other organs caused by various reasons; Complicated with systemic infection, immune system deficiency, or unstable vital signs; Patients with UPJO other than pyeloureteral junction stenoses, such as concomitant renal or ureteral calculi, ectopic vascular compression, iatrogenic stenosis, duplex kidney or ureter, ureteral polyps, and other causes, were all removed. Children with a BMI of 24 or more. According to the inclusion criteria, 34 children (28 males and 6 females) were selected as research objects. According to the inclusion and exclusion criteria, 34 children (28 boys and 6 girls) were selected as the study subjects.

Surgical intervention is the main treatment of UPJO caused by ureteropelvic junction stricture. At present, dismembered pyeloplasty is the first choice for treatment. The principle of operation showed that the effect of this method on relieving obstruction was immediate, and timely intervention could reduce the degree of renal impairment or further prevent its aggravation^[4]^. In this study all the children underwent dismembered pyeloplasty, open surgery, or laparoscopic surgery performed based on the age of the children. The children over 5 years old were treated with laparoscopy, and the other children were treated with small incision minimally invasive open surgery.

The surgical indications are: The diagnosis was confirmed and UPJO was present. Auxiliary examination showed that obstruction had impaired renal function^[5]^. Imaging examination showed obvious obstruction.

Sampling and surgical procedure: All children were reexamined by urinary B-ultrasound and urine routine after admission. Before the operation, nuclear magnetic resonance hydrography, intravenous pyelography, biochemical, and other auxiliary examinations were completed, and diagnosis and evaluation were performed. Surgical contraindications were excluded and preoperative preparation was perfected.

After the anesthesia takes effect in children with open surgery, the urine bag is clamped, the healthy side position is adjusted, and disinfection and sheet spreading are often prescribed. An oblique incision of approximately 3-4cm in length was made in the 11th intercostal space on the affected side. Each layer was incised sequentially to the retroperitoneal space, and the perirenal fascia was bluntly separated, the middle and lower part of the kidney, the renal pelvis, and the upper ureter were exposed free, dilated, and separated renal pelvis and distinct narrow genocidal junction were seen. Mark the highest and lowest two sites of the renal pelvis. The proximal ureter was released by mobilizing the surrounding tissue, and the lower pole of the kidney was located by searching the lower renal calyx. When the ureteropelvic junction and proximal ureter were fully exposed, obliquely cut the renal pelvis, and the stenotic ureter was resected at the same time. The normal ureter facing the renal pelvis was gently lifted and incised 1.0-1.5 cm along the wedge, carefully judged until the normal ureteral tissue appeared. Normal ureteral tissue can be clearly distinguished from the diseased segment under the naked eye. The lesion segments were not only small and tough but also had internal white scar feeling, dry and narrow thin cords. Therefore in this study, the lesion segment was taken as the center, the upper edge of the renal pelvis was taken as the control group, and the lower edge of the ureteral stenosis segment was also taken to study whether there was a difference of pathological changes between them. During the operation, the lesion segment was not only small and tough but also the internal white scar felt, dry, narrow and thin cord-like. The stricture and dysplastic ureteropelvic junction and its surrounding tissues were resected and sent for pathological examination within half an hour. The extra-renal pelvis was smoothed cut, and the lower pole of the kidney and the lowest point of the ureterotomy were fixed with intermittent para-position suture. The renal pelvis was reconstructed by continuous suturing after 4-6 stitches^[6]^. The double J ureteral tube was inserted, and the renal pelvis was reset after the suturing was completed. One negative pressure drainage tube was disposed of in the renal fossa, and the incision was closed by suturing layer by layer. If laparoscopic pyeloplasty is selected, except for the steps and some details such as positioning, placing the stamp card, establishing pneumoperitoneum, and closing the instrument entrance after an operation, the other steps are the same as above.

Pathological examination methods: The moderate and severe hydronephrosis caused by UPJ stenosis in UPJO was considered to be caused by abnormal development of the ureteropelvic junction tissue. The surgical intervention will remove the stenotic lesion segment, anastomose the normal tissue of the upper renal pelvis and the lower ureter, and restore the peristaltic function so that urine drainage is unobstructed. Reducing or eliminating hydronephrosis achieves the purpose of protecting kidney function. Therefore, from the diseased segment of the UPJ as the center, the 1.5 cm segment of the renal pelvis was taken along the upper edge and upward (as the control group), and the 1.0 cm ureteral stricture tissue was taken along the lower edge and downward. To explore the etiological mechanism by comparing the microscopic pathological characteristics of the diseased segment and the normal segment, all the specimens obtained were fixed with a tissue fixator. Tissues were embedded in paraffin and sectioned, each piece was approximately 4 microns thick, hematoxylin and eosin were used for HE staining, Masson trichrome staining, and immunohistochemistry to observe and evaluate the distribution of collagen fibers and ICC cells^[7]^.

### 2.2 Data collection

HE staining and Masson trichrome staining (by Solarbio) follow the reagent instructions. And the immunohistochemical method (The antibody reagents used were from Elabscience) is xylene and gradient alcohol were used for dewaxing and drying. An appropriate amount of antigen repair solution EDTA was prepared and heated with distilled water in a water bath for 20 minutes to repair the antigen and the antigen was naturally cooled to room temperature. Each section was delimited around the tissue with a hydrophobic pen and rinsed with PBS solution, and 10mg of lyophilized HRP powder was dissolved in PBS and added to the section to block. After the primary antibody was diluted, an appropriate amount of it was added on top of the slices, covered with the wet box, and then translated to 4 degrees Celsius in the refrigerator overnight. The next day, after washing with PBS, an appropriate amount of secondary antibody was added and placed for 40 minutes. The color development was performed at the same time right after it was ready. The cells were counterstained using the prepared hematoxylin for 30 seconds, rinsed in PBS, and returned to blue in distilled water. It was immersed in absolute ethanol for dehydration and allowed to dry. Each piece was sealed by dropping an appropriate amount of neutral gum.

### 2.3 Statistical analyses

The target we observe is the muscle shape of different parts was observed. The observable indicators are MC ratio (the muscle to collagen ratio was analyzed by an independent observer through image color analysis. After cutting and staining the target area of the specimen, the online color extraction tool ImageJ Fiji2 was used to calculate the area of color difference in the slices and quantify the MC ratio. The optic red area corresponds to muscle, and the blue area corresponds to collagen. The ratio was calculated separately and converted into a single number, and the ratio of the two was taken as the MC ratio, e.g., 60%/40%, Z1.5.) and the number of positive cells in IHC was counted (2 observers, double-blind evaluation, counted the target cells in each upper left, lower left, middle, upper right, lower right and 9 fields between them. Because CD117 is expressed in both mast cells and ICC cells, nonspecific staining should be excluded when counting.

The statistical treatment we use is statistical software SPSSAU to analyze the data, the measurement data of MC ratio (muscle to collagen ratio) and the number of ICC cells in different tissue parts. Multiple sample Friedman test and paired T test were used for data analysis.

## RESULTS

### 1. Pathological changes of the lesions and IHC

**FigureI :**
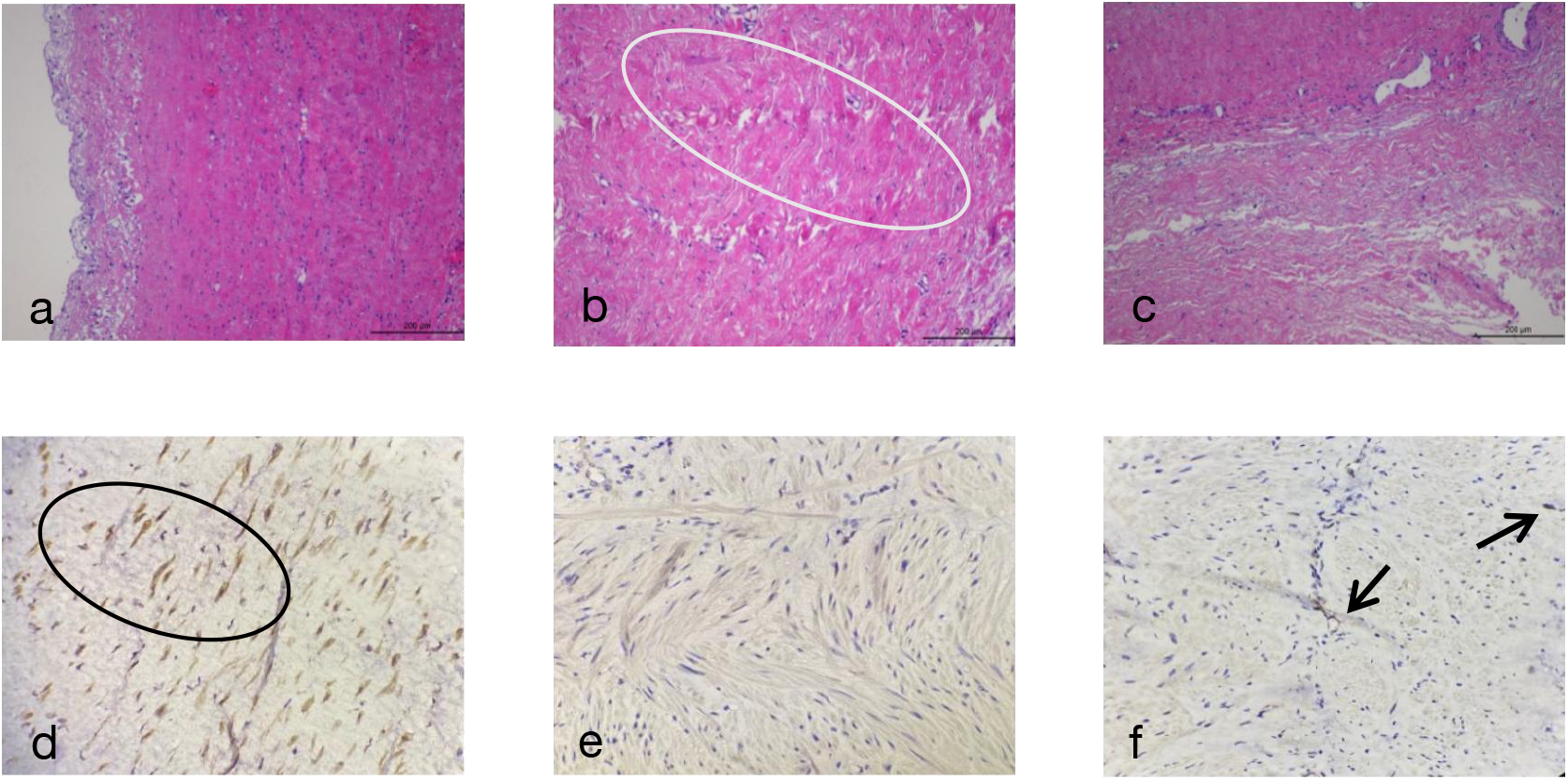
**Fig.a** The muscle layer of the renal pelvis segment is arranged in a regular pattern (×100). The smooth muscle of the tube wall shows the inner longitudinal and outer ring. The urothelium and the underlying lamina propria can be seen, and the layers are clear and consistent. **Fig.b** The muscle layer of the pathological part of the renal pelvis and ureter connection is disordered and increased by hyperemia, and a large number of collagen fibers can be seen in the muscle layer, as shown in the white circle (× 100). The muscle layer is hypertrophic, disorderly arranged and not dense, and a large amount of hyperplasia collagen fibers can be seen in the muscle layer, inflammatory cell infiltration was also observed. **Fig.c** The ureteral stricture showed chronic inflammation of the mucosa, and the interstitial fibrous tissue showed obvious proliferation, with a small amount of chronic inflammatory cell infiltration. **Fig.d** More DAB chromogenic cells can be found in the renal pelvis; **Fig.e** In the lesion segment of UPJ, the number of chromogenic cells is small or even absent, the connective tissue is hyperplasia, and the arrangement of histiocytes is disordered; **Fig.f** The ureteral stricture segment also showed fibrous hyperplasia, which was lighter than the lesion segment, with a few scattered chromogenic cells.

### 2. MC ratio at different sites

As shown in Table 1, the proportion of muscle and collagen in specimens of different parts was obtained by image analysis software. It was obtained by Friedman test of multiple paired samples.Significant differences were found between the RP segment vs. the UPJ segment and between the RP segment vs. the US segment, and no significant differences were found between the UPJ segment and the US segment.

**Table 1.** MC ratio (muscle to collagen ratio) in renal pelvic segment, UPJ segment and stenosis segment.

| | Median ± SD | Statistical Magnitude | P value |
| --- | --- | --- | --- |
| RP | 0.5 ± 0.351 | | |
| UPJ | 0.06 ± 0.024 | 54.692 | 0.001 |
| US | 0.09 ± 0.045 | | |

### 3. Masson trichrome stain

**FigureII:**
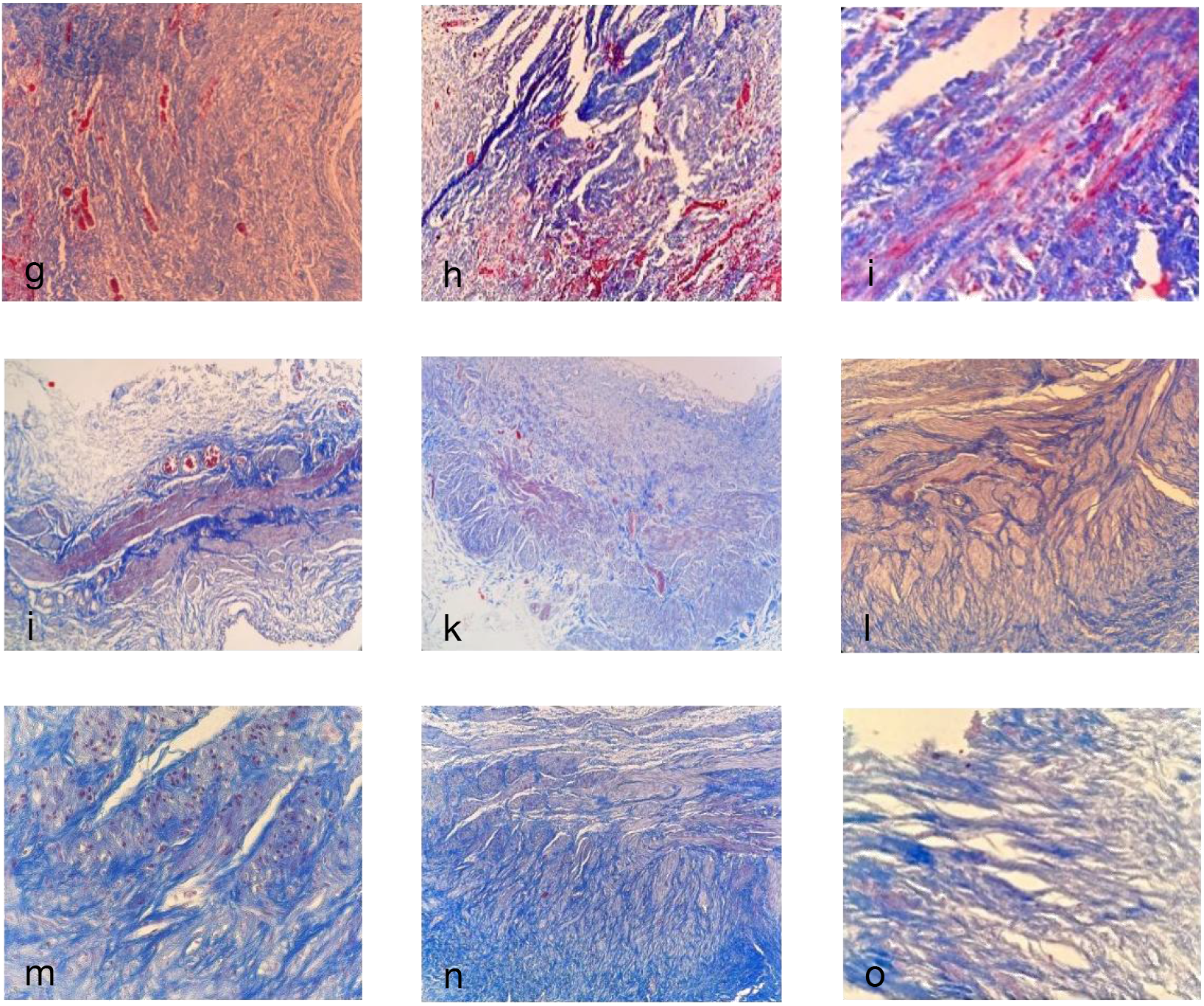
**Fig. (g, h, i)** The distribution of muscle and fibrous tissue in the renal pelvis segment is shown, with red representing muscle and blue representing fibrous tissue. Fig.f shows that the muscle fibers are arranged in regular shape, which corresponds to the muscle distribution of the renal pelvis segment in HE staining. **Fig. (j, k, l)** The tissue in the lesion section of UPJ was shown, and the muscle distribution is significantly reduced and the arrangement is disordered with varying degrees of connective tissue hyperplasia, and the normal muscle tissue has disappeared. **Fig. (m, n, o)** The distribution of muscle tissue is less than that of the renal pelvis segment, but compared with the lesion segment, muscle tissue is still visible, and there is also hyperplasia of connective tissue around. The fibrous hyperplasia is heavier than the lesion segment and the renal pelvis segment.

### 4. The number of ICC and CVF in different parts were compared

### 5.

**Figure III :**
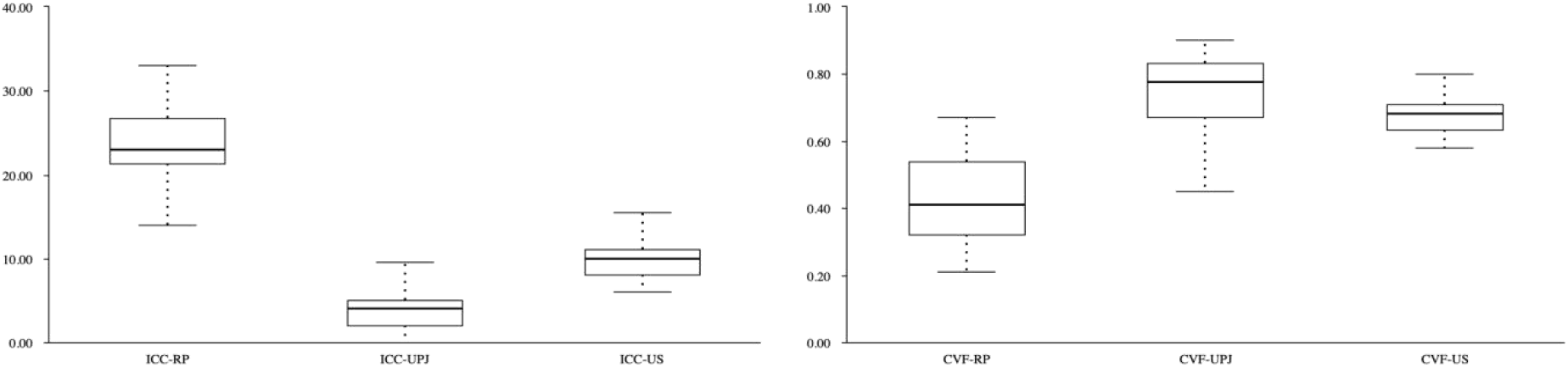
Box plots of ICC and CVF at different sites.

As shown in Table 2 and Box plot : Median and standard deviation of interstitial cells of Cajal in nine different high power fields in visible renal pelvis segment, UPJ lesion segment, and ureteral stricture segment. The results were significantly different after SPSSPPRO using Freidman test and Nemenyi test. The number of ICC cells in the renal pelvis was the most, while the distribution of UPJ lesion was the least or even absent. Collagen volume fraction CVF derived from collagen distribution in Masson trichrome stained specimens obtained by Fiji image analysis software. Using t-test analysis, the collagen volume fraction in the renal pelvis segment, UPJ lesion segment and ureteral stenosis segment were respectively different. The distribution of collagen fibers in the diseased and stenosis segments of UPJ was less different than that in the other two segments. The collagen volume fraction was the highest in the lesion segment of UPJ, and the collagen fibers were significantly proliferated.

**Table 2.** Comparison of ICC cell number and collagen volume fraction (CVF) between different sites.

| Segments |  | Median ± SD |  | Statistic |  |  |  |
| --- | --- | --- | --- | --- | --- | --- | --- |
|  |  |  |  | al<br>Magnitu<br>de | T value | P value | Cohen's d |
| ICC(<br>num<br>ber) | RP vs. UPJ | 23 ± 4.729 | 19 ± 2.234 | 11.49 | -19.913 | 0.001 | 5.118 |
|  | RP vs. US | 23 ± 4.729 | 13 ± 2.665 | 6.002 | 16.376 | 0.001 | 3.700 |
|  | UPJ vs. US | 4 ± 2.496 | 6 ± 0.431 | 5.488 | -9.751 | 0.001 | 2.556 |
| CVF<br>(%<br>) | RP vs. UPJ | 0.41 ± 0.131 | 0.365 ± 0.021 | 9.861 | -11.097 | 0.001 | 2.825 |
|  | RP vs. US | 0.41 ± 0.131 | 0.27 ± 0.079 | 7.117 | -10.321 | 0.001 | 2.679 |
|  | UPJ vs. US | 0.775 ± 0.11 | 0.095 ± 0.058 | 2.744 | 3.233 | 0.003 | 0.874 |

## Conclusion

In children with hydronephrosis caused by ureteropelvic junction stenosis, the number of interstitial cells of Cajal and the degree of collagen fiber proliferation in the diseased segment of UPJ were significantly different from those in the upper edge of the renal pelvis and the lower edge of the ureter stenosis. However, the ratio of muscle to collagen was the highest in the renal pelvis and was different from that in the UPJ and stenosis segments, there was no significant difference in MC ratio between the UPJ segment and the stenotic segment. The principle of the operation is to connect the upper and lower normal ureteral tissue. It is suggested that although clinicians are concerned about the length of the ureter and to prevent excessive tension at the suture, they should also resect all the ureteral stricture at the lower edge of the ureteropelvic junction as much as possible to ensure the removal of abnormal tissue in the ureter as much as possible and relieve the mechanical obstruction.

## Discussion

Ureteropelvic junction obstruction (UPJO) is the most common disease leading to congenital hydronephrosis in children and its etiology is complex, among which UPJ stenosis is the most common cause. Among them, the stenosis of UPJ is the most common cause of disease. From the perspective of histopathology, Some studies have found that the diseased segment of the ureteropelvic junction usually has hypertrophy and disarrangement of smooth muscle, and there is a large number of collagen fiber hyperplasia between the muscles. The interstitial cells of Cajal with pacing function distributed in the urinary tract were widely reduced in number, sparsely distributed, or even absent in the diseased segment. There were significant differences compared with adjacent normal tissues such as renal pelvis and proximal ureter tissue, On the one hand, it was verified that the lack of ICCs in the lesion segment resulted in ineffective down-flow of urinary tract peristalsis, obstruction and hydronephrosis[12], there is also a chronic inflammatory change in the mucosa with infiltration of inflammatory cells, accompanied by disarrangement of muscle fibers. Various kinds of collagen fibers proliferated in the muscle and other pathological changes. As the child grows, the kidneys continue to produce urine. Without timely intervention at the right time, the hydronephrosis will further expand and cause long-term compression of the nephron due to the limited space in the trunk. There is a great risk of varying degrees of renal function damage in the long term, which seriously endangers the health of children. For children with clear symptoms and auxiliary examination to verify the existence of exact obstruction, only surgical intervention can relieve the obstruction at present. However, due to the characteristics of pyeloureteroplasty itself, there is still the possibility of long-term anastomotic stenosis, secondary stone and other complications. Long-term follow-up and relevant examinations are needed to evaluate the recovery of children.

Among the known major causes of UPJ stenosis, changes in interstitial cells of Cajal are considered to be of great importance[13], the number of ICCs at the ureteropelvic junction is closely related to the occurrence of UPJO in children with congenital hydronephrosis. However, it remains to be elucidated why the number of ICCs at the ureteropelvic junction decreases. Pathological examination of the specimens showed inflammatory cell infiltration in most cases, early obstructive symptoms may be related to the presence or recurrence of urinary tract infection and the degree of inflammation. Inflammatory response may promote scar hyperplasia in the lesion segment, thereby promoting further stenosis at the stenotic site. From the perspective of clinicopathology, this study compared the pathological changes in different anatomical parts of the surgically resected tissue in the same child to verify and explore possible etiological direction. In this study, 34 surgical specimens were selected and divided into three parts according to their anatomical structures. They were renal pelvis segment, ureteropelvic junction lesion segment, ureteral stenosis segment. HE staining, Masson trichrome staining and immunohistochemical staining were performed to explore the number and distribution of interstitial cells of Cajal in different parts, the arrangement of muscle, and the distribution ratio of muscle tissue to collagen fibers. To test the etiological hypothesis of UPJO caused by UPJ stenosis. According to the experimental results, it can be concluded that the muscle tissue of the diseased segment of UPJ is disordered and the collagen fibers between the muscle are proliferated. The number distribution of interstitial cells of Cajal was lower than that of the two. As the normal control group, there were more muscle tissue in the renal pelvis, and the shape was orderly arranged in the inner longitudinal and outer rings, and there was no defect in the urothelium. The number of interstitial cells in Cajal with pacemaker function was the highest, the number of ICCs in the ureteral stenosis segment was more than that in the lesion segment and renal pelvis segment. It is consistent with previous studies that the ureter develops from the ureteral bud from the center to both ends during embryonic development, and the occurrence of lesion segments reflects that the mechanism of such diseases may be the outcome of developmental arrest.

It should be noted that ICCs need to be differentiated from mast cells when counting by immunohistochemistry. Is due to the fact that both interstitial cells of Cajal and mast cells express CD117 (which is c-kit positive). It is feasible to distinguish by their morphological features[9–3]. Interstitial cells of Cajal are generally located at the inner edge of the circular myometrium, parallel to the muscle fibers, they had spindle cell bodies, thin and few cytoplasm, large nuclei, generally oval, and usually 2 dendrites. Mast cells had a round central nucleus and were commonly found in the submucosa, mucosa, and between the mucosa, while c-kit positive ICCs were generally only found in the muscular layer. Therefore, the number of ICCs should be calculated by subtracting the number of mast cells from the number of CD117-positive cells in the specified field of view.

The results showed that a large number of c-kit positive ICCs were observed in normal renal pelvis tissues. In the lesion segment of UPJ, ICCs were sparse or absent, and the density of ICCs in samples from different sites was statistically different. In addition, there was a band of brown positive color distribution arranged like “rain shape” in the renal pelvis specimen, which was considered to be the long tail Tp of Tc cells mentioned in previous studies and should not be counted as interstitial cells of Cajal.

The sample size of this study is relatively small, and although the selected samples meet the inclusion and exclusion criteria, there are still biases caused by different duration of disease and inconsistent degrees of obstruction. In the future, it is hoped that this study can be continued in a multi-center, large number of samples, and more accurate stratified sampling by age and quantitative assessment of obstruction severity. We hope that more powerful evidence will emerge to prove this etiological hypothesis.

The wide range of deficits in the children receiving the intervention studied here, however, reliable studies have shown that age-related changes in the expression of CD117-positive ICCs do not alter the distribution of c-kit-positive ICCs in UPJO[11], so the reliability of the conclusion is increased.

In conclusion, after comparing the pathological changes in different anatomical parts of the same specimen, it is inferred that the cause of UPJ stenosis is related to the decrease in the number of interstitial cells of Cajal, the disarrangement of muscle tissue and the proliferation of collagen fibers in the diseased segment. It is suggested that the etiology of UPJO can be further explored to the inducing factors of proliferation and decline of interstitial cells of Cajal, the fibrosis process of lesion tissue and the correlation, also whether it is affected by the inflammatory response between them.

## Notes

### Competing Interest Statement

The authors have declared no competing interest.

## REFERENCES

1. Yhoshu, E., Menon, P., Rao, K., & Bhattacharya, A. (2022). Outcome Analysis of Reduction and Nonreduction Dismembered Pyeloplasty in Ureteropelvic Junction Obstruction: A Randomized, Prospective, Comparative Study. Journal of Indian Association of Pediatric Surgeons, 27(1), 25–31.

2. Babu, R., Vittalraj, P., Sundaram, S., & Shalini, S. (2019). Pathological changes in ureterovesical and ureteropelvic junction obstruction explained by fetal ureter histology. Journal of pediatric urology, 15(3), 240.e1–240.e7.

3. Mohammadjafari, H., Rafiei, A., Kosaryan, M., Yeganeh, Y., & Hosseinimehr, S. J. (2014). Determination of the severity of ureteropelvic junction obstruction using urinary epidermal growth factor and kidney injury molecule 1 levels. Biomarkers in medicine, 8(10), 1199–1206.

4. Abbas, T., Elifranji, M., Al-Salihi, M., Ahmad, J., Vallasciani, S., Elkadhi, A., Özcan, C., Burgu, B., Akinci, A., Alnaimi, A., & Salle, J. (2022). Functional recoverability post-pyeloplasty in children with ureteropelvic junction obstruction and poorly functioning kidneys: Systematic review. Journal of pediatric urology, S1477-5131(22)00313-8. Advance online publication.

5. Vemulakonda V. M. (2021). Ureteropelvic junction obstruction: diagnosis and management. Current opinion in pediatrics, 33(2), 227–234.

6. Wishahi, M., Mehena, A. A., Elganzoury, H., Badawy, M. H., Hafiz, E., & El-Leithy, T. (2021). Telocyte and Cajal cell distribution in renal pelvis, ureteropelvic junction (UPJ), and proximal ureter in normal upper urinary tract and UPJ obstruction: reappraisal of the aetiology of UPJ obstruction. Folia morphologica, 80(4), 850–856.

7. Liu, G., Liu, X., & Yang, Y. (2022). Comparative transcriptome analysis of miRNA in hydronephrosis male children caused by ureteropelvic junction obstruction with or without renal functional injury. PeerJ, 10, e12962.

8. Huang, Z. P., Qiu, H., Yang, Y., Zhang, L., Yang, B., Lin, M. J., & Yu, B. P. (2016). The Role of Interstitial Cells of Cajal in Acute Cholecystitis in Guinea Pig Gallbladder. Cellular physiology and biochemistry : international journal of experimental cellular physiology, biochemistry, and pharmacology, 38(5), 1775–1784.

9. Mut, T., Acar, Ö., Oktar, T., Kılıçaslan, I., Esen, T., Ander, H., & Ziylan, O. (2016). Intraoperative inspection of the ureteropelvic junction during pyeloplasty is not sufficient to distinguish between extrinsic and intrinsic causes of obstruction: Correlation with histological analysis. Journal of pediatric urology, 12(4), 223.e1–223.e2236.

10. Issi, O., Deliktas, H., Gedik, A., Ozekinci, S., Bircan, M. K., & Sahin, H. (2015). Does the histopathologic pattern of the ureteropelvic junction affect the outcome of pyeloplasty. Urology journal, 12(1), 2028–2031.

